# Bringing Down Cancer Aircraft: Searching For Essential Hypomutated Proteins In Skin Melanoma

**DOI:** 10.1101/020396

**Authors:** Mikhail Pyatnitskiy, Dmitriy Karpov, Ekaterina Poverennaya, Andrey Lisitsa, Sergei Moshkovskii

**Affiliations:** Institute of Biomedical Chemistry, 119121, Pogodinskaya str, 10, Moscow, Russia; ZAO Personal Biomedicine, 129164, Prospekt Mira, 124, 17, Moscow, Russia; Pirogov Russian National Research Medical University, 117997, Ostrovityanova str,1, Moscow, Russia; Engelhardt Institute of Molecular Biology, 119991, Vavilova str, 32, Moscow, Russia

**Keywords:** cancer genome, negative selection, Cancer Genome Atlas, cancer essential protein, high-impact mutations, anti-tumor therapies

## Abstract

Abstract

**Background:** We propose an approach to detection of essential proteins required for cancer cell survival. Gene is considered essential if mutation with high functional impact upon function of encoded protein causes death of cancer cell. We draw an analogy between essential cancer proteins and well-known Abraham Wald’s work on estimating the plane critical areas using data on survivability of aircraft encountering enemy fire. Wald reasoned that parts hit least on the returned planes are critical and should be protected more. Similarly we propose that genes essential for tumor cell should carry less high-impact mutations in cancer compared to polymorphisms found in normal cells.

**Results:** We used data on mutations from the Cancer Genome Atlas and polymorphisms found in healthy humans (from 1000 Genomes Project) to predict 91 protein-coding genes essential for melanoma. These genes were selected according to several criteria including negative selection, expression in melanocytes and decrease in the proportion of high-impact mutations in cancer compared with normal cells.

Gene ontology analysis revealed enrichment of essential proteins related to membrane and cell periphery. We speculate that this could be a sign of immune system-driven negative selection of cancer neo-antigens. Another finding is overrepresentation of semaphorin receptors, which can mediate distinctive signaling cascades and are involved in various aspects of tumor development. Cytokine receptors CCR5 and CXCR1 were also identified as cancer essential proteins and this is confirmed by other studies.

**Conclusions:** Overall our goal was to illustrate the idea of detecting proteins whose sequence integrity and functioning is important for cancer cell survival. Hopefully, this prediction of essential cancer proteins may point to new targets for anti-tumor therapies.

## Background

Recent progress in genome sequencing in frame of large cancer studies conferred a new vision of tumor as a result of mutator phenotype [1]. Instead of earlier point of view that cancer cell has mutations affecting only specific oncogenes or tumor suppressor genes it became clear that its genome is literally filled up with various somatic abberations [2]. A strategy of survival and evolution of the cancer cell clone is to change its genome quickly using damage or modulation of DNA repair systems [3]. Some mutations are drivers of the cancer process and occur in the cancer-related genes. However, most mutations occurring throughout the whole genome are not relevant to the tumor progression and represent passenger mutations. They do not help cancer cells to survive and may experience a negative selection [4].

In recent works where data from cancer genome sequencing were analyzed a pivotal attention is paid to the identification of genes significantly mutated in cancer compared to germline genome. A statistical study made on thousands of samples has reported more than 200 potential cancer driver genes [5]. The acquisition of specific mutations in these genes is the driving force of malignant transformation.

Our study represents an alternative approach to analysis of cancer genome data. The idea is inspired by well-known fact from the history of applied statistics, Abraham Wald’s aircraft problem [6]. Wald proposed to search for aircraft vulnerability zones by estimation of the bullet-free patches on airplanes which returned to the airbase from the combat flight. Indeed, it is the bullet-free areas on the machine surface are essential for the aircraft performance. If bullets hit those areas, then machines crashed and data on aircraft vulnerability became unobservable (Fig.1, left panel). So the bullet-free zones on the returned aircrafts are essential for plane functioning and hence needed more armor.

**Figure 1.**
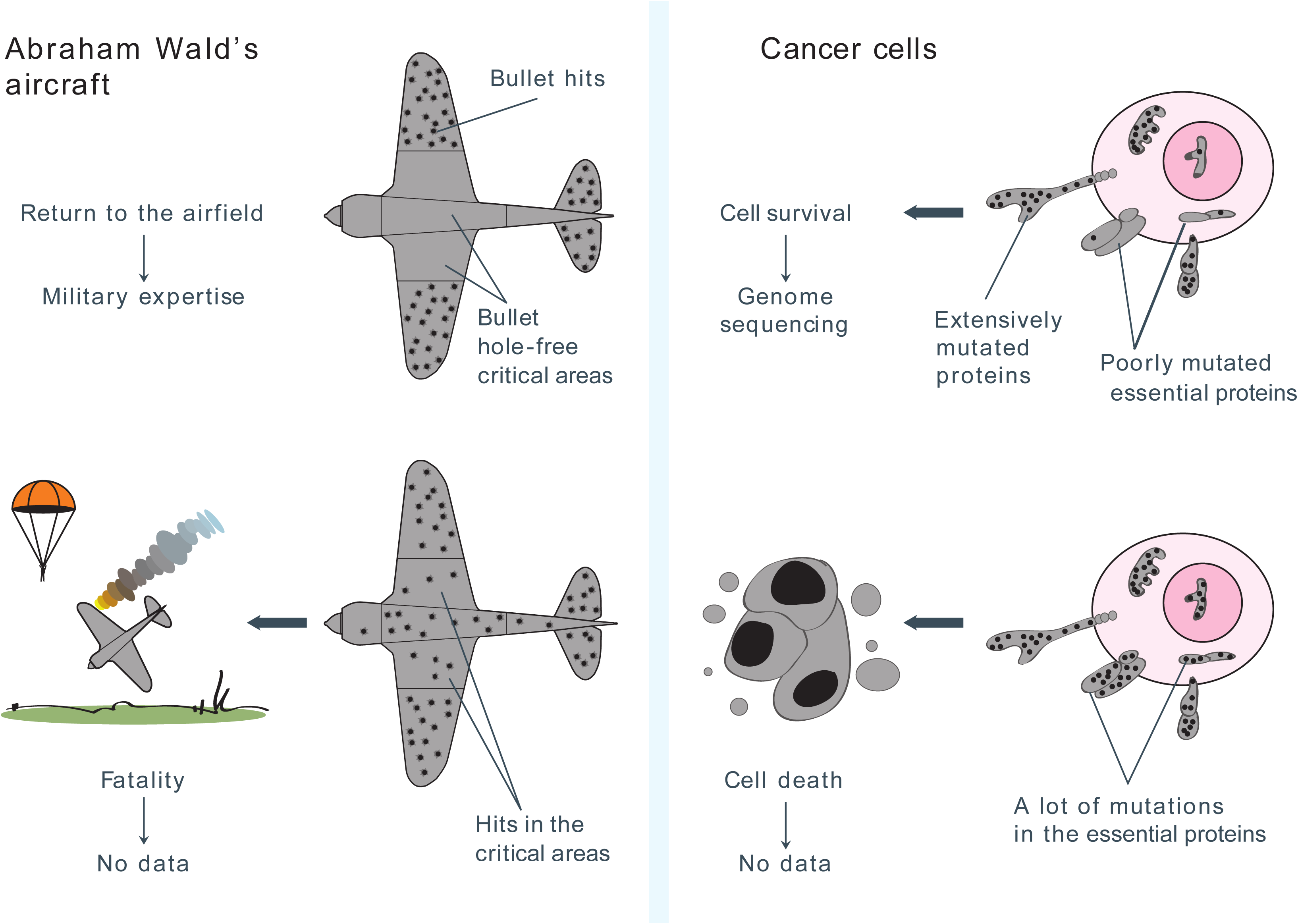
Analogy between bullet-free plane critical areas and hypomutated proteins essential for cancer. Undamaged areas on the returned planes are critical for the aircraft performance. Similarly we propose that proteins with reduced number of deleterious somatic mutations compared to germline are essential for cancer cell survival.

We have envisaged that the airplane surface is a cancer genome and bullets are somatic mutations with high impact upon protein function. The sites with decreased number of such variants may be essential for cancer survival, since the cells with mutations in these sites die and their genomes are not sequenced by cancer projects (Fig.1, right panel).

Expectedly, we were not alone in such way of thinking. In the beginning of cancer genomics, it was suggested to search homozygous DNA deletions as immutable features of cancer cells [7]. In his recent work, Polak et al. analyzed hypomutated sites in cancer genome and found them in accessible regulatory DNA due to enhanced repair in these sites [8].

However, we focused not on the cancer genome, but rather on cancer proteome. We searched for the hypomutation phenomenon from the viewpoint of protein functioning. Our goal was to identify proteins which are depleted by functionally important amino acid changes in cancer. These proteins are conserved and therefore are essential for cell survival during cancer evolution. As a reference we used polymorphisms reported in 1000 Genomes Project [9]. In fact, we have compared evolution of protein sequences during development of cancer with such evolution during 200,000 years of development of humanity from its last bottleneck [10]. We believe that our approach is able to detect proteins whose sequence integrity and functioning is more important for cancer cells rather than for normal tissues. Hopefully, these predicted cancer-essential proteins may represent vulnerability zones for tumor cells and hence serve as new targets for anti-cancer drugs.

## Methods

Cancer mutation data were downloaded from Synapse website [11], accession number syn1729383. These data initially were obtained via The Cancer Genome Atlas [12] and were reprocessed to filter out false positives and germline variants, details can be found in [13]. Skin cutaneous melanoma dataset (SKCM) had the highest number of variants compared to other datasets, total 181175 variants. Data on germline polymorphisms were downloaded from 1000 Genomes (1KG) website [9] and annotated using Annovar software [14]. Final table contained 425069 variants located within coding regions. Genes with less than 11 variants either in SKCM or in 1KG data were removed from subsequent analysis.

For each human protein-coding gene the ratio of the rates of non-synonymous and synonymous substitutions (dN/dS) was calculated as in [15] using SKCM data on somatic mutations. Genes with dN/dS value greater than 0.25 (i.e. experiencing either neutral or positive selection) were removed.

Gene expression data were obtained via TCGA website and contained information about 20531 genes expressed in 374 melanoma samples. Gene expression profiles that had absolute expression levels in the lowest 20% of the dataset were removed.

Functional impact of non-synonymous single nucleotide variants in both SKCM and 1KG data was assessed using dbNSFP resource [16]. This database compiles results from ten prediction algorithms (including SIFT, Polyphen2 and MutationAssessor) into two ensemble scores, MetaSVM and MetaLR. Larger value of score indicates that the variant is more likely to be damaging. We considered variant to have high impact on protein functioning if either MetaSVM or MetaLR value was greater than 70-th percentile of corresponding score. Somatic mutation was also considered to have high functional impact if it belonged to one of the following classes: insertion/deletion, stop loss, stop gain, or splice site mutation. For each gene we calculated fraction of mutations with high functional impact *f*_1KG_ and *f*_SKCM_ as *f = #(high func. mutations) / #(total mutations)*

Functional geneset enrichment analysis via hypergeometric test was performed using WebGestalt software [17]. As a reference set we supplied list of 16172 genes with average SKCM expression greater than 20-th percentile. Correction for multiple testing was done using Benjamini & Hochberg method.

A list of interactions for selected proteins was downloaded from STRING database v. 9.1 [18] using Python script. STRING collects various associations between proteins, such as structural predictions, textmining, pathway analysis and experimental results from other web resources, to form its aggregative score [19]. Interactions with the score greater than 0.9 were used for analysis. Network visualization and analysis was performed via Cytoscape [20].

## Results and discussion

The purpose of our analysis was to identify cancer proteins under negative selection i.e. depleted with functionally important amino acid changes. These conserved proteins may be essential for cell survival during cancer evolution and hence can be regarded as possible targets for anticancer therapy (Fig 1).

For our analysis we chose melanoma somatic mutation dataset available from TCGA website. Our approach is assumed to be more applicable to tumors triggered by point mutations rather than by copy-number alterations [21], such as lung tumors or melanomas. Those former cancers provide much more statistics on protein sequences to make the prediction robust and melanoma is characterized by the highest somatic mutation frequency as compared to other cancers [22]. Also melanoma dataset contained the highest number of variants compared to other types of cancer, total 181,175 variants.

### Defining a subset of skin melanoma proteins under negative selection

In order to find protein-coding genes experiencing negative selection during cancer evolution we used *d*_N_/*d*_S_ as an indicator of selective pressure [23]. The same logic was accepted by Ostrow et al [15] when they studied positive selection in cancer genomes.

We calculated dN/dS ratio for all human protein coding genes using SKCM data. Genes carrying less than 11 variants were excluded from analysis in order to obtain more robust estimates of dN/dS ratio. Gene was considered to be under negative selection if corresponding dN/dS value was smaller than predefined threshold. In attempt to avoid threshold choice by arbitrary decision we plotted genome-wide distribution of dN/dS values for three cancer types with largest number of point mutations, i.e. two lung cancers and skin melanoma (Fig. 2). As can be seen from the histograms, local minima can observed at dN/dS value about 0.25. We used this threshold as a first filter to select essential cancer genes which were supposed to be under negative selection.

**Figure 2.**
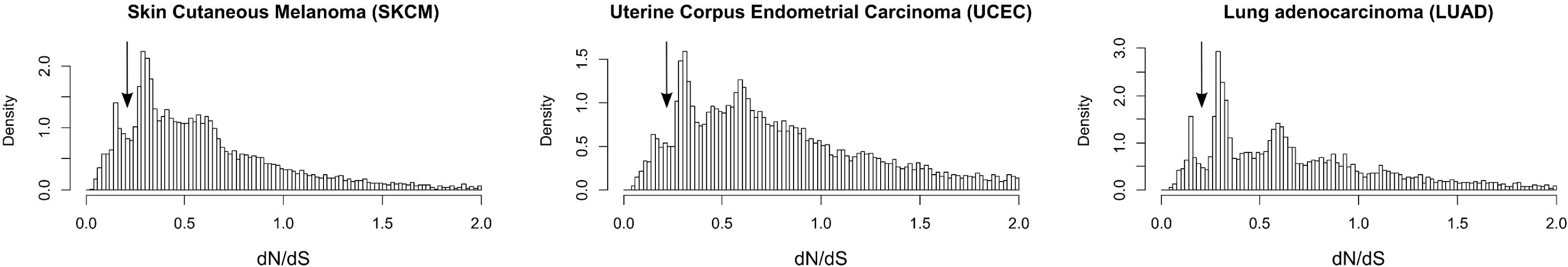
Genome-wide distribution of dN/dS ratio for three cancer types. Skin melanoma, uterine endometrial carcinoma and lung adenocarcinoma contained most of the data on cancer mutations. Local minima are reached when dN/dS is approximately 0.25 (denoted by arrows). We considered proteins with dN/dS below this threshold to experience negative selection.

Our next step was to remove genes which are not expressed in melanoma cells and hence cannot be recognized as essential for this type of cancer. We calculated average expression for each gene across 374 cancer samples and removed genes with expression levels in the lowest 20%.

Then we turned to the evaluation of genetic variants observed in SKCM and 1KG data. We classified mutations into two classes: those with high impact on the protein function and other non-significant mutations, including synonymous variations. Functional effect of non-synonymous substitutions was predicted via dbNSFP database [16] (see Methods). Indels, stop gains and losses and changes occurring within splice sites were also classified as mutations with high functional impact [see Additional file 1].

Finally we selected genes which are depleted by functionally important variants in cancer as compared to normal tissues, i.e. genes where fraction of mutations with high functional impact *f* for 1KG data was greater than the same fraction for cancer data, *f*_1KG_ > *f*_SKCM_. After all steps we obtained 91 protein-coding genes designated hereinafter as “essential cancer proteins”.

### Skin melanoma essential protein subset is significantly enriched by plasma membrane proteins: a possible link to immunobiology

By manual looking through the list, one could mark some distinct categories among those proteins (Table 1, Additional file 1). Most represented categories included membrane transport proteins, such as ion channels and solute carriers, neural proteins of various functions, cell adhesion molecules, etc. In order to describe the resulting list more formally we performed functional enrichment analysis using WebGestalt website [17].

**Table 1.**
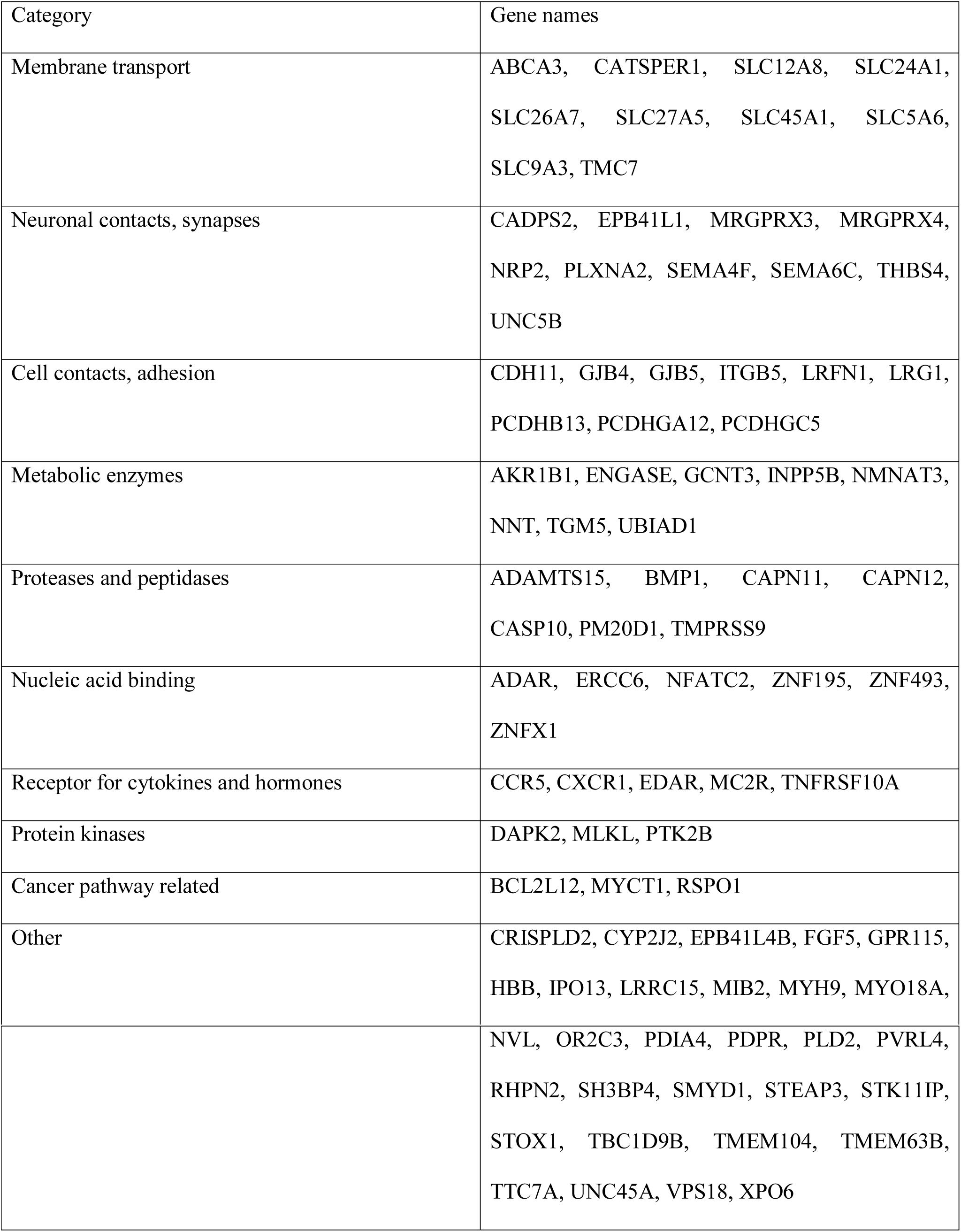
List of protein-coding genes with amino acid sequences under negative selection in skin melanoma genomes (essential cancer proteins). Genes were filtered as described here. Categories were defined by manual biocuration.

Enrichment of 91 essential protein subset by Gene Ontology [24] categories had shown a significant trend towards membrane proteins, specifically, proteins of plasma membrane and cell periphery (Table 2). Obviously, such general categories cannot fully decipher possible molecular pathways or cell proliferation mechanisms. However, there may be a mechanistic interpretation of the overrepresented plasma membrane proteins. It is widely recognized that cancers escape from host immunity through evolution of cancer clones [25]. We have hypothesized that the conservation of plasma membrane and cell periphery protein sequence in melanoma could be a result of such immune escape. More specifically, high impact mutations in cell periphery and plasma membrane proteins may lead to formation of major histocompatibility complex (MHC) II-dependent neo-antigens [26]. MHC II-restricted protein epitopes reactive to T-helper lymphocyte subpopulation, as widely known, are formed by digestion of phagocytized extracellular proteins and cell periphery protein accompanying internalized membrane parts. The fact, that such mutations are depleted in cell surface and periphery proteins, reflects escape of melanoma cells from CD4^+^-T-cell mediated immunity. Recently it has been reported, that adoptive immunity induced against a T-helper-1 (MHC II-restricted) neo-antigen epitope provided tumor regression in a patient with metastatic cholangiocarcinoma [27]. Significant response of CD4+-T-cells against personalized tumor neo-epitopes was also found in patients with metastatic melanoma [28]. Thus, we may observe the enrichment by cell periphery proteins in target subset as a result of purifying selection against formation of neo-antigen T-helper epitopes. When a plasma membrane protein is extensively mutated in a cancer cell, it is digested after vesicle internalization and its mutant peptides bind MHC II as neo-epitopes which are not recognized by immune system as self-epitopes. Having such neo-epitopes expressed, a cancer cell cannot avoid the immune surveillance and is eliminated (Fig. 3). Notably, despite MHC II itself is expressed in limited cell types, such as professional antigen-presenting cells, melanoma cells are reported to express various types of the receptor [29].

**Figure 3.**
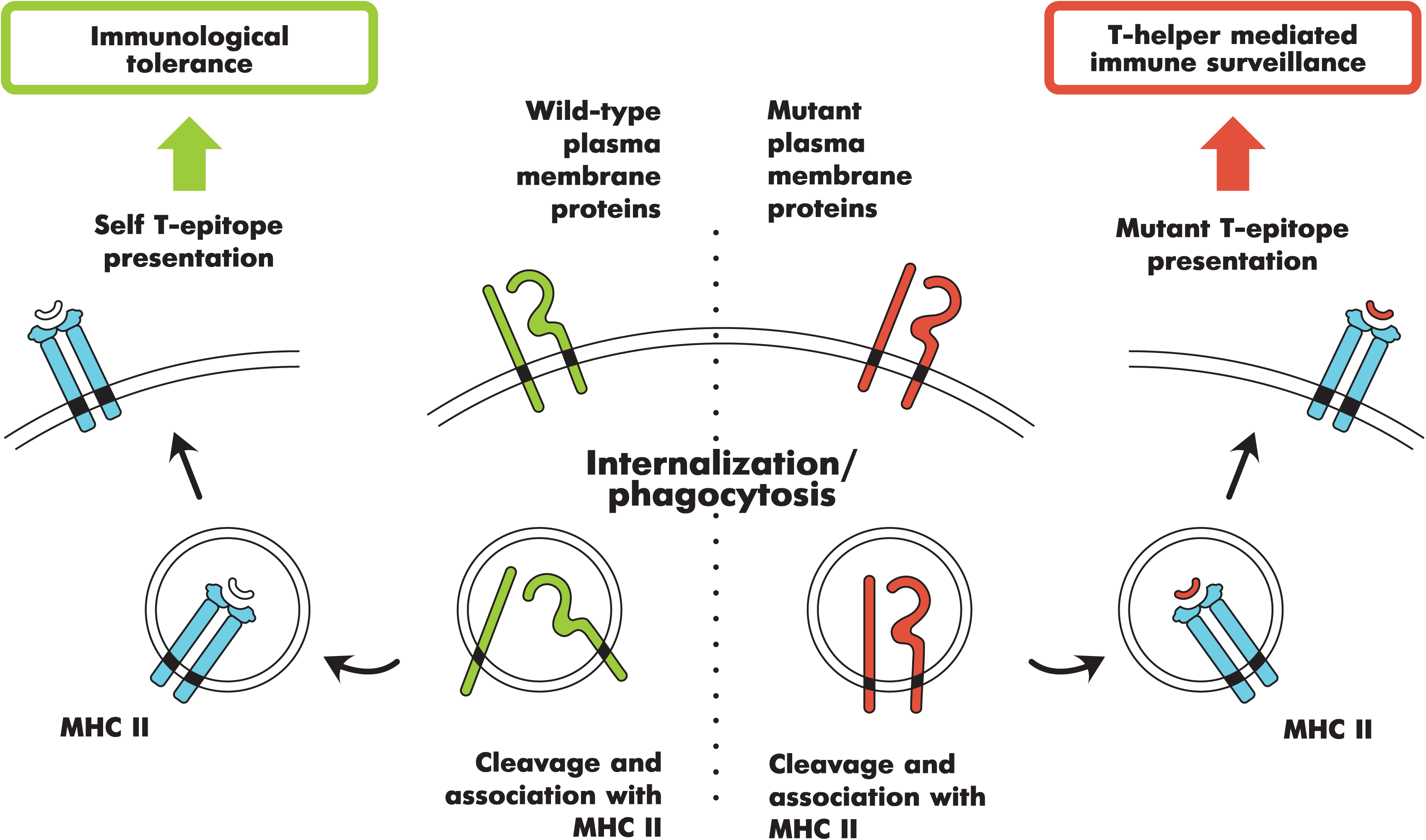
Hypothetical scheme for negative selection against neoantigens derived from cell surface proteins. Preferential involvement of surface proteins to MHC-restricted antigen presentation is known in the art [26]. Cancer cells exposing MHC-II epitopes with mutated antigens are more likely to be eliminated by T-cell mediated immune surveillance.

**Table 2.** Enrichment of the essential cancer protein subset by Gene Ontology, KEGG [51] and PharmGKB drug target [52] categories. P-values were adjusted using Benjamini-Hochberg correction for multiple comparisons.

| Class | Name | adjusted<br>p-value | Genes |
| --- | --- | --- | --- |
| GO cellular component | membrane part | 0.0002 | ABCA3, CATSPER1, CCR5, CDH11, CXCR1, CYP2J2, EDAR, EPB41L1, EPB41L4B, GCNT3, GJB4, GJB5, GPR115, INPP5B, ITGB5, LRFN1, LRRC15, MC2R, MRGPRX3, MRGPRX4, MYH9, NNT, NRP2, OR2C3, PCDHB13, PCDHGA12, PCDHGC5, PLD2, PLXNA2, PTK2B, PVRL4, SEMA4F, SEMA6C, SH3BP4, SLC12A8, SLC24A1, SLC26A7, SLC27A5, SLC45A1, SLC5A6, SLC9A3, STEAP3, TBC1D9B, TMC7, TMEM104, TMEM63B, TMPRSS9, TNFRSF10A, UBIAD1, UNC5B, VPS18 |
| GO cellular component | integral to membrane | 0.0004 | ABCA3, CATSPER1, CCR5, CDH11, CXCR1, EDAR, GCNT3, GJB4, GJB5, GPR115, INPP5B, ITGB5, LRFN1, LRRC15, MC2R, MRGPRX3, MRGPRX4, MYH9, NNT, NRP2, OR2C3, PCDHB13, PCDHGA12, PCDHGC5, PLXNA2, PVRL4, SEMA4F, SEMA6C, SLC12A8, SLC24A1, SLC26A7, SLC27A5, SLC45A1, SLC5A6, SLC9A3, STEAP3, TBC1D9B, TMC7, TMEM104, TMEM63B, TMPRSS9, TNFRSF10A, UBIAD1, UNC5B |
| GO cellular component | intrinsic to membrane | 0.0004 | ABCA3, CATSPER1, CCR5, CDH11, CXCR1, EDAR, GCNT3, GJB4, GJB5, GPR115, INPP5B, ITGB5, |
|  |  |  | LRFN1, LRRC15, MC2R, MRGPRX3, MRGPRX4, MYH9, NNT, NRP2, OR2C3, PCDHB13, PCDHGA12, PCDHGC5, PLXNA2, PVRL4, SEMA4F, SEMA6C, SLC12A8, SLC24A1, SLC26A7, SLC27A5, SLC45A1, SLC5A6, SLC9A3, STEAP3, TBC1D9B, TMC7, TMEM104, TMEM63B, TMPRSS9, TNFRSF10A, UBIAD1, UNC5B |
| GO cellular component | plasma membrane | 0.0004 | ABCA3, CASP10, CATSPER1, CCR5, CDH11, XCR1, EDAR, EPB41L1, GJB4, GJB5, GPR115, ITGB5, LRFN1, MC2R, MRGPRX3, MRGPRX4, MYH9, NFATC2, NRP2, OR2C3, PCDHB13, PCDHGA12, PCDHGC5, PLD2, PLXNA2, PTK2B, PVRL4, SEMA4F, SEMA6C, SLC24A1, SLC26A7, SLC27A5, SLC5A6, SLC9A3, STEAP3, TMPRSS9, UNC5B, XPO6 |
| GO cellular component | cell periphery | 0.0006 | ABCA3, CASP10, CATSPER1, CCR5, CDH11, CXCR1, EDAR, EPB41L1, GJB4, GJB5, GPR115, ITGB5, LRFN1, MC2R, MRGPRX3, MRGPRX4, MYH9, NFATC2, NRP2, OR2C3, PCDHB13, PCDHGA12, PCDHGC5, PLD2, PLXNA2, PTK2B, PVRL4, SEMA4F, SEMA6C, SLC24A1, SLC26A7, SLC27A5, SLC5A6, SLC9A3, STEAP3, TMPRSS9, UNC5B, XPO6 |
| GO cellular component | membrane | 0.0019 | ABCA3, CADPS2, CASP10, CATSPER1, CCR5, CDH11, CXCR1, CYP2J2, EDAR, EPB41L1, EPB41L4B, GCNT3, GJB4, GJB5, GPR115, INPP5B, |
|  |  |  | ITGB5, LRFN1, LRG1, LRRC15, MC2R, MRGPRX3, MRGPRX4, MYH9, NFATC2, NNT, NRP2, OR2C3, PCDHB13, PCDHGA12, PCDHGC5, PLD2, PLXNA2, PTK2B, PVRL4, SEMA4F, SEMA6C, SH3BP4, SLC12A8, SLC24A1, SLC26A7, SLC27A5, SLC45A1, SLC5A6, SLC9A3, STEAP3, TBC1D9B, TMC7, TMEM104, TMEM63B, TMPRSS9, TNFRSF10A, UBIAD1, UNC5B, VPS18, XPO6 |
| GO cellular component | semaphorin receptor complex | 0.0112 | NRP2, PLXNA2 |
| KEGG | axon guidance | 0.0126 | NFATC2, PLXNA2, SEMA4F, SEMA6C, UNC5B |
| Drug associated genes | enfuvirtide | 0.0153 | CCR5, PDIA4 |
| Drug associated genes | ribavirin | 0.0153 | ADAR, CCR5, NMNAT3 |
| Drug associated genes | naloxone | 0.0153 | MRGPRX3, MRGPRX4 |
| GO molecular function | receptor activity | 0.0197 | CCR5, CXCR1, EDAR, GPR115, ITGB5, MC2R, MRGPRX3, MRGPRX4, NRP2, OR2C3, PLXNA2, SEMA4F, SEMA6C, TNFRSF10A, UNC5B |
| GO molecular function | transmembrane signaling receptor activity | 0.0197 | CCR5, CXCR1, EDAR, GPR115, MC2R, MRGPRX3, MRGPRX4, NRP2, OR2C3, PLXNA2, TNFRSF10A, UNC5B. |
| GO molecular<br>function | secondary active<br>transmembrane<br>transporter<br>activity | 0.0197 | SLC12A8, SLC24A1, SLC26A7, SLC45A1, SLC5A6,<br>SLC9A3 |
| GO molecular<br>function | active<br>transmembrane<br>transporter<br>activity | 0.0266 | ABCA3, SLC12A8, SLC24A1, SLC26A7, SLC45A1,<br>SLC5A6, SLC9A3 |
| GO cellular<br>component | brush border<br>membrane | 0.0286 | PLD2, SLC5A6, SLC9A3 |
| GO molecular<br>function | semaphorin<br>receptor activity | 0.029 | NRP2, PLXNA2 |
| GO molecular<br>function | signaling<br>receptor activity | 0.029 | CCR5, CXCR1, EDAR, GPR115, MC2R, MRGPRX3,<br>MRGPRX4, OR2C3, PLXNA2, TNFRSF10A, UNC5B,<br>NRP2 |
| GO molecular<br>function | transporter<br>activity | 0.029 | ABCA3, CATSPER1, GJB4, HBB, IPO13, NNT,<br>SLC12A8, SLC24A1, SLC26A7, SLC27A5, SLC45A1,<br>SLC5A6, SLC9A3, XPO6 |
| GO molecular<br>function | calcium-<br>dependent<br>cysteine-type<br>endopeptidase<br>activity | 0.029 | CAPN11, CAPN12 |
| GO molecular<br>function | chemokine<br>binding | 0.029 | CCR5, CXCR1 |
| GO molecular | G-protein | 0.029 | CCR5, CXCR1, GPR115, MC2R, MRGPRX3, |
| function | coupled receptor activity |  | MRGPRX4, OR2C3 |
| GO molecular function | symporter activity | 0.029 | SLC12A8, SLC24A1, SLC45A1, SLC5A6 |
| GO molecular function | antiporter activity | 0.0303 | SLC24A1, SLC26A7, SLC9A3 |
| GO molecular function | signal transducer activity | 0.0303 | CCR5, CXCR1, EDAR, GPR115, MC2R, MIB2, MRGPRX3, MRGPRX4, NRP2, OR2C3, PLXNA2, PTK2B, TNFRSF10A, UNC5B |
| GO molecular function | molecular transducer activity | 0.0303 | CCR5, CXCR1, EDAR, GPR115, MC2R, MIB2, MRGPRX3, MRGPRX4, NRP2, OR2C3, PLXNA2, PTK2B, TNFRSF10A, UNC5B |
| GO cellular component | connexon complex | 0.0381 | GJB4, GJB5 |
| GO molecular function | cation:cation antiporter activity | 0.0384 | SLC24A1, SLC9A3 |
| GO molecular function | cytokine binding | 0.0384 | CCR, CXCR1, NRP2 |
| GO molecular function | heparin binding | 0.0396 | CRISPLD2, NRP2, RSPO1, THBS4 |
| GO molecular function | cysteine-type endopeptidase activity | 0.0435 | CAPN11, CAPN12, CASP10 |

Immunoediting of the cancer genome against antigen formation as described recently [30] is in a good correspondence with our hypothesis.

Certainly, we do not intend that MHC II directed negative selection is the only factor forming hypomutated species of the subset. It should be considered as one of many factors influencing the overall purifying selection during cancer evolution.

### CCR5 and CXCR1 chemokine receptors as essential melanoma proteins

One of the proteins in our subset is a C-C chemokine receptor type 5 (*CCR5*), a chemokine receptor which was extensively studied due to its ability to provide HIV fusion with the target cell. Moreover, a deletion of this protein known as CCR5-Δ32 protects its carrier against selected strains of the virus [31]. Thus, in case of changed amino acid sequence of the receptor it does not serve as a cofactor for the virus particle. Drugs blocking CCR5, e.g. maraviroc, are considered as a therapy of choice for the treatment of HIV infection.

At the same time, a role of CCR5 in cancer has been widely discussed. It is known that this receptor is often overexpressed in some cancer types [32]. CCR5 expression has been induced after oncogenic transformation of cells [33]. Evidences are accumulating that this receptor’s signaling through its ligand, CCL5, provides cancer progression, e.g. metastasis. CCR5 antagonist drugs have been shown to block metastasis of basal breast cancer [34] and Src-induced prostate cancer [35] in mice. It was proposed to use well-tolerable CCR5 inhibitor drugs, which are already approved for use in HIV infection, in clinical trials for metastatic cancers [34]. An observational trial that studied effect of maraviroc on metastatic colorectal cancer had been completed, but no results have been reported yet [35].

As for melanoma, a recent paper has shown that growth and metastasis of this tumor was inhibited in CCR5^-/-^ mutant mice [36]. The authors have proposed that CCR5/CCL5 signaling negatively affect CD8 T-lymphocyte effects. Thus, melanoma cells expressing functional CCR5 may thereby contribute to the immune escape.

The fact, that C-C chemokine receptor type 5 is one of essential cancer proteins, positively illustrates applicability of our approach. This receptor is a druggable cancer target, at least, for adjuvant therapy.

Another important cytokine receptor in our list is CXCR1. It binds interleukin-8 protein (CXCL8) which is a major chemotaxis agent in innate immunity. Notably, CXCR1 and above-mentioned CCR5 use the same signaling pathways and are closely related in their function [37]. As for the latter, CXCR1 is known to be used by melanoma cells for outgrowth and metastasis [38]. Recently, it was shown that elevated expression of this receptor is correlated with tumor malignancy [39]. Most likely, CCR5 and CXCR1 are used by melanoma cells to survive in inflammatory environment.

### Semaphorin pathway is potentially important for melanoma survival

Another promising enrichment among the essential cancer proteins is related to the semaphorin receptor complex and axon guidance pathway (Table 2). Semaphorins are involved in cell guidance during development and in adult tissues [40]. These proteins were recently shown to function as suppressing and promoting agents in various cancer types via transactivation of receptor tyrosine kinases [40]. Notably, both types of semaphorin-binding receptors, plexins and neuropilins, are represented in our list and may be involved in cancer cell survival [42]. Plexin A2 (*PLXNA2*) protected cancer cells from death after human papillomavirus infection [43]. For neuropilin-2 (*NRP2*), there is an evidence of its role in malignant melanoma where its expression was correlated with tumor progression [44]. These results provide a background for further experimentation to discover cancer drug targets among semaphorin complex proteins [40].

### Protein interactions between essential melanoma proteins

In order to understand how predicted essential melanoma proteins cooperate with each other, we analyzed corresponding protein-protein interactions using STRING database [18]. We have selected only high-confident interactions with score greater than 0.9. Interacting preys were reported for 46 of 91 genes of interest (Table 3). Most of protein interactions are known for cytokine receptors CCR5 (115 preys) and CXCR1 (77 preys), of them about 70 interactions involve common partners. Another pair of interactors which are characterized by many common partners consists of caspase-10 (CASP10) and TRAIL-R1 death receptor (TNFRSF10A).

**Table 3.** Protein-protein interactions: the number of preys reported for essential melanoma proteins as baits according to STRING database [18]. Only highly reliable interactions with score greater than 0.9 were considered.

| Gene name(s) | Number of<br>interacting preys |
| --- | --- |
| CCR5 | 115 |
| CXCR1 | 77 |
| PLD2 | 45 |
| NFATC2 | 44 |
| INPP5B | 36 |
| PTK2B | 35 |
| CYP2J2 | 31 |
| MC2R | 27 |
| ERCC6 | 26 |
| ADAR | 25 |
| ITGB5 | 24 |
| AKR1B1,<br>CASP10, PLXNA2 | 15 |
| CDH11, OR2C3 | 12 |
| GCNT3,<br>NMNAT3 | 11 |
| TNFRSF10A | 10 |
| UNC5B | 8 |
| BMP1, HBB,<br>MYH9 | 7 |
| NRP2, SLC27A5 | 6 |
| EPB41L1, FGF5,<br>IPO13, SLC9A3 | 4 |
| PVRL4, RSPO1,<br>STEAP3, THBS4,<br>VPS18 | 3 |
| ABCA3,<br>CATSPER1, NNT,<br>PDIA4, STK11IP,<br>UBIAD1 | 2 |
| EDAR, MIB2,<br>MLKL, MYCT1,<br>MYO18A, UNC45A | 1 |

The interaction map of proteins hypomutated in melanoma (Fig. 4) can be easily subdivided to several main networks. First, three G-coupled receptors CCR5, CXCR1 and MCR2 form a triangle of many interactions. A role of these receptors in melanoma where they probably promote cancer cell survival and metastasis is partly described above.

**Figure 4.**
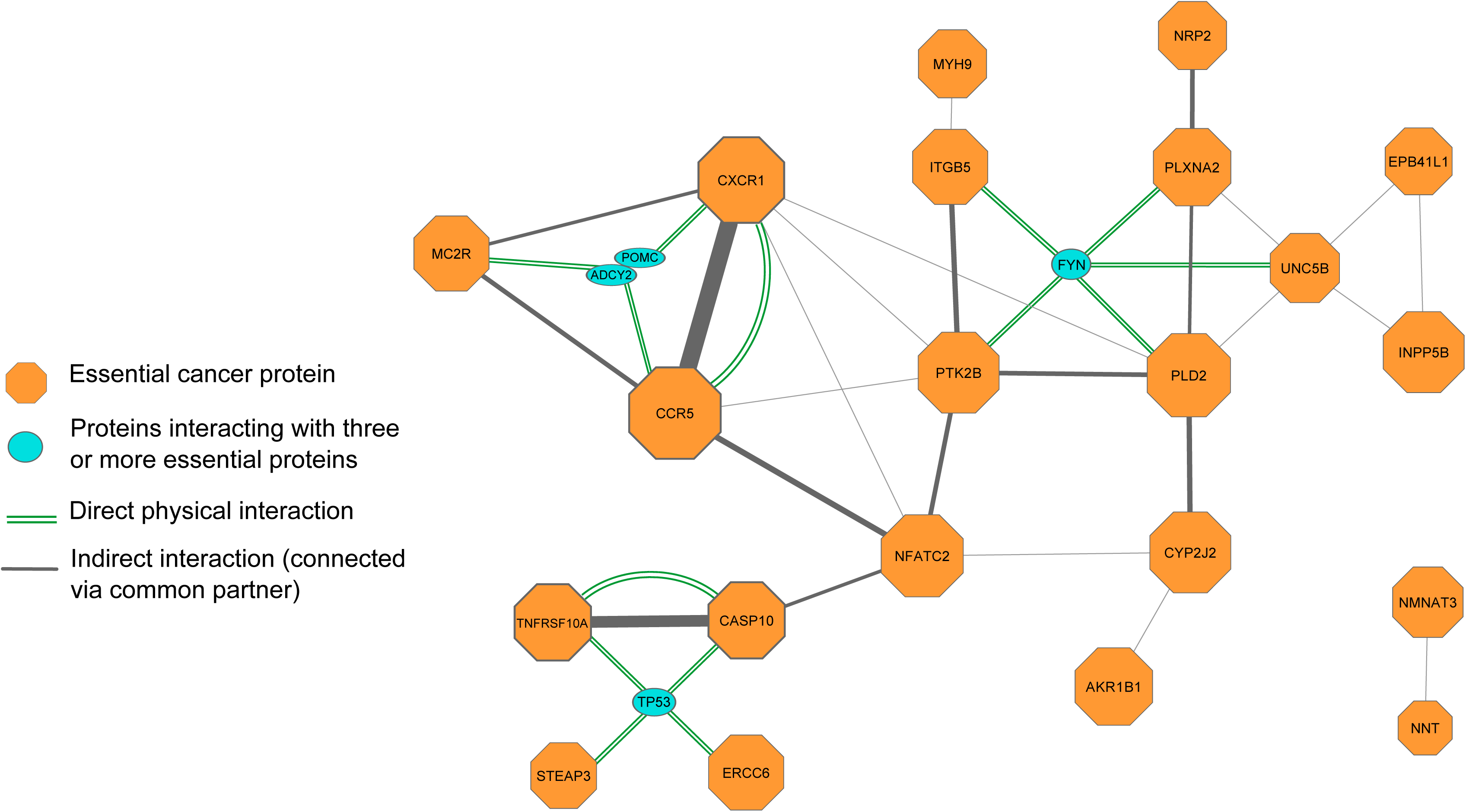
Interaction map for essential melanoma proteins. The map was built using STRING database with high-confidence interaction score threshold 0.9. Size of the octagon is proportional to the number of partners of corresponding protein. Line width reflects the number of common partners between two baits. Doubled green lines denote known direct physical protein-protein interactions. Prey proteins interacting with three or more baits are shown as blue ovals: FYN, TP53, ADCY2 and POMC.

The largest network component between integrin ITGB5, protein kinase PTK2B, semaphorin receptors (PLXNA2) and phospholipase D2 (PLD2) is defined by their interactions with FYN protein (Fig. 4). The latter is a well-known protein kinase from the Src family and is involved in the series of cellular functions, such as T-cell immunity, axon guidance and is also considered as a potential proto-oncogene [45]. Little is known about its function in melanoma and there is some evidence that Fyn cannot be a medication target in advanced melanoma. In particular, when it was inhibited by saracatinib drug along with other Src kinases, no benefit was observed in advanced melanoma [46]. However, based on our data, more attention should be paid to study a role of Fyn in melanoma separately from other Src kinases, such as Src itself, Yes, etc.

Unexpectedly, amongst hypomutated proteins, some partners of TP53 tumor suppressor were found. They include, inter alia, physically interacting proteins, caspase-10 (CASP10) and TRAIL-R1 death receptor (TNFRSF10A). In contrast to these results, components of TRAIL apoptosis pathway are considered to have antitumor effect, when active [47]. This contradiction awaits further deciphering.

## Conclusions

The cancer genome concept developed over the past decade has literally revolutionized our understanding of cancer molecular biology [48]. Results of cancer genome projects are especially promising for evidence-based and personalized medicine. Mutational landscape of any cancer may disclose driver genes which provide tumor development and progression [5]. At the same time, not all driver proteins may serve as targets for existing or possible drug therapies. Many cancers are primarily caused by tumor suppressors which are irreversibly inactivated by gene mutations. In these cases, drug therapy represents a big challenge [49] due to difficulties in complementation of the lost function, e.g. by gene therapy. Therefore a search for new drug targets is a primary task of post-cancer-genome studies.

Positive selection regulates, at the level of corresponding genes, amino acid sequences of driver cancer proteins during clone competition accompanying tumorigenesis. Necessity of oncogenes and tumor suppressors to be mutated in most cancers inspires researchers to implement scoring systems to select hypermutated driver genes [5]. In contrast to these useful efforts, instead we focused on proteins whose corresponding genes are hypomutated in tumor, i.e. experience negative selection during cancer evolution. We believe that these proteins may provide additional drug targets especially in cases where cancer drivers represent suppressors functionally destroyed by mutations.

For the analysis we have chosen skin melanoma exome dataset available from TCGA project [50], because this cancer is characterized by the highest level of point mutations.

With our approach aggregating evolutional dN/dS parameter as a measure of negative selection, gene expression and functional impact of amino acid changes, we have predicted 91 protein-coding genes to be essential in melanoma. Despite the list of proteins was a mixture of entities of various structures and functions, we detected several significant enrichments which let us generate explanatory hypotheses. For example, the list contained increased number of cell surface and cell periphery proteins. In our opinion, it could be a sign of immune system-driven negative selection of cancer neo-antigens [28]. Furthermore, some examples of hypomutated proteins represent known cancer-related proteins, such as cytokine receptors CCR5 and CXCR1 [37].

Obviously, our data is preliminary. This work is mostly intended to illustrate a general idea of defining essential cancer proteome. Results may become more reliable when more individual cancer genomes would be accumulated. However, even at the present form, our list of predicted essential cancer proteins provides a background for targeted experimentation with tumor cell survival by blocking the protein candidates *in vitro* and *in vivo*.

## Availability of supporting data

The data set supporting the results of this article is included within the article.

### List of abbreviations

SKCM: skin cutaneous melanoma
TCGA: The Cancer Genome Atlas
1KG: 1000 Genomes Project
HIV: human immunodeficiency virus

## Authors' contributions

MP carried out computational experiments and helped to draft the manuscript. DK participated in experiments. EP performed the interactomics analysis. AL helped to draft the manuscript and interpreted the data. SM drafted the manuscript and designed the experiments. All authors read and approved the final manuscript.

## Acknowledgments

The work was supported by the Russian Scientific Fund, grant # 14-15-00395 to SM.

## Additional files

Additional file 1. Format: tab-separated text

List of protein-coding genes with amino acid sequences under negative selection in skin melanoma genomes (“essential” cancer proteins).

Column description:

Gene, EntrezGene: gene identifiers according to HGNC and NCBI Gene Description gene function description
skcm.dnds: dN/dS ratio calculated using SKCM data on somatic mutations. quantile.expression percentile of gene expression (TCGA SKCM data) norm.neutral number of synonymous SNVs (1KG data)
norm.high: number of mutations with high impact functional impact (1KG data) skcm.neutral number of synonymous SNVs (TCGA SKCM data)
skcm.high: number of mutations with high impact functional impact (TCGA SKCM data)

